# Suppression and facilitation of motion perception in humans

**DOI:** 10.1101/465807

**Authors:** Tzvetomir Tzvetanov

## Abstract

In a study that investigated the putative neuronal origin of suppression and facilitation in human motion perception Schallmo et al. (2018) used various techniques to investigate how motion perception is shaped by excitatory and inhibitory interactions within hMT+ or earlier areas. They further proposed that neuronal normalization is a sufficient account of the behavioural results, discounting accepted and precise neuronal mechanisms of excitation or inhibition. In this Research Advance, it is shown (1) that once the full computational model that predicts the psychophysical results is defined, it is not “divisive normalization” but actually excitatory and inhibitory processes that are the neuronal mechanisms shaping facilitation and suppression in the behavioural domain, then (2) that the experimental design they used allows a *quantitative* comparison and usage of such “contrast–size tuning” data.

## Introduction

Changes in perception are interesting scientific tools in the search of how and where is visual perception created in our brain (Spillmann & Dresp, 1995; Spillmann & Ehrenstein, 1996; Spillmann & Werner, 1996; Eagleman, 2001; Albright & Stoner, 2002; Born & Bradley, 2005). Human motion perception is known to exhibit various astonishing effects, among which very known in the public domain are motion illusions (for some examples among many websites, see https://michaelbach.de/ot/index.html), with even simple motion stimuli providing sometimes counter-intuitive results.

Concerning the percept of a single motion direction, that is, of an object or a collection of objects (as rain drops) moving in a single unidirectional motion, it is considered to be created from the interaction between excitatory, pooling information across space and directions, and inhibitory, suppressing information across space and directions, interactions between motion sensitive neurons. For example, the percept of random-dot kinematograms/patterns (RDK/RDPs) was reported to be strongly shaped by the context in which the target motion was presented (Tynan & Sekuler, 1975; Levinson & Sekuler, 1976; Marshak & Sekuler, 1979; Chang & Julesz, 1984; Watamaniuk, Sekuler & Williams, 1989; Nawrot & Sekuler, 1990). Limiting the topic to the spatial spread of the interactions, two interesting studies by two Japanese scientists (Murakami & Shimojo, 1993; Murakami & Shimojo, 1996) showed that size tuning of these centre-surround contextual interactions in motion perception seems to be eccentricity invariant. That is, when rescaling the measures for different eccentricities all results naturally fell on a single curve. This somewhat recalls the size tuning characteristics of receptive fields and their magnification with visual eccentricity. The authors proposed a simple spatial centre-surround model of receptive fields for explaining their observations.

Yet in another type of psychophysical probes, D. Tadin and colleagues demonstrated that perception of simple moving grating stimuli (Tadin, Lappin, Gilroy & Blake, 2003), or a type of RDK (Tadin & Lappin, 2005), have very peculiar perceptual results with respect to size and contrast of the stimuli (see Figure 1A). Globally large and low contrast stimuli needed much less presentation time for being perceived than large and high contrast stimuli, while the opposite was observed for small stimuli. That is, there is an inverted effect on perception as a function of contrast for small and large stimulus sizes (Figure 1A). These observations were also attributed to the contrast and size tuning properties of motion tuned neuronal populations.

**Figure 1:**
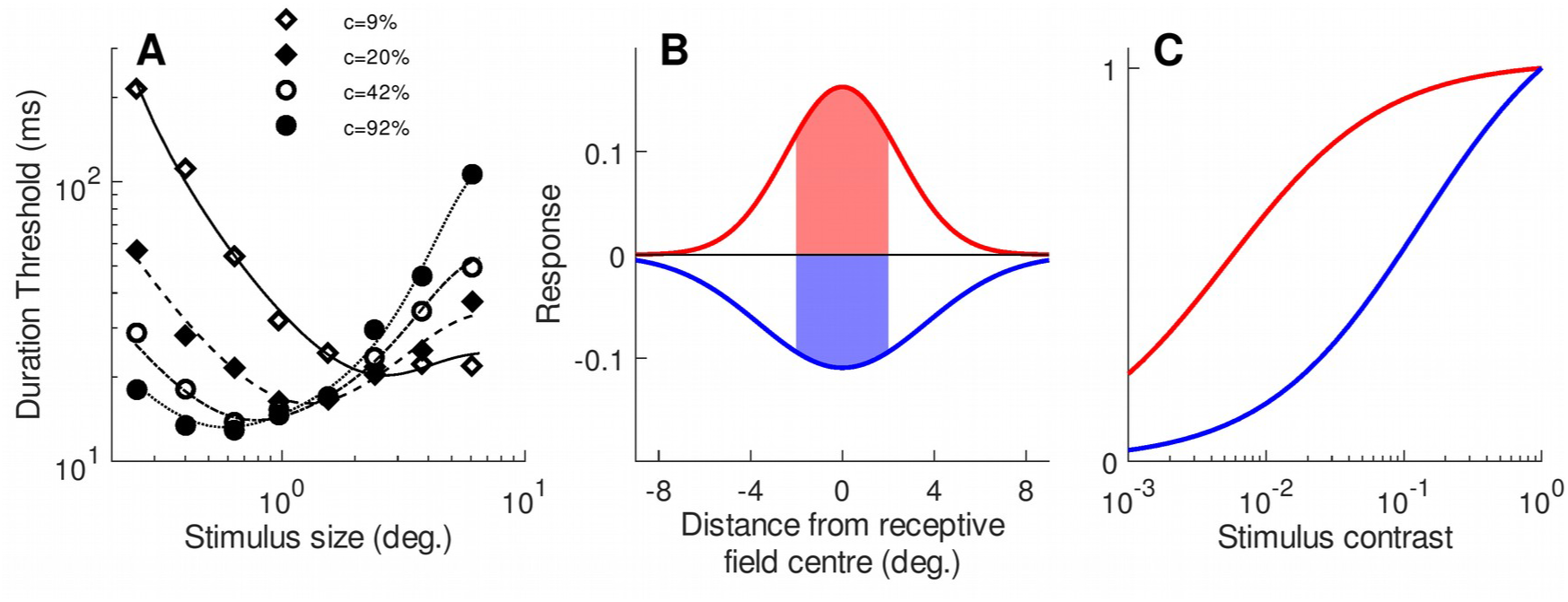
(A) Typical report of effects of stimulus contrast and size on duration thresholds (data extracted and replotted from Figure 1A in Tadin and Lappin (2005)). Duration Threshold is the necessary stimulus presentation time in order to correctly perceive its direction of motion (see **Psychophysics** for details). Curves are model fits (see **Modelling** and Equation 15). (B-C) Spatial and contrast tuning characteristics of the receptive field components. (B) illustration of spatial receptive field components – excitatory (red) and inhibitory (blue) Gaussian profiles with their total response to receptive-centred stimulus of diameter 4 degrees (red and blue shaded surfaces). (C) Contrast response functions of excitatory (red) and inhibitory (blue) components of the receptive field.

This tuning has a simple form with excitatory and inhibitory components (Figure 1B,C). It has a visuospatial receptive field structure with a relatively small excitatory centre but spatially larger inhibitory surround (panel B) and contrast responses such that the excitatory component is activated much rapidly at low contrasts (panel C). The final neuronal activity is a combination of the responses of these two drives. It is their precise shape and method to combine them that gives the particular property of the neurons. If one creates stimuli that somewhat match the preferred characteristics of these neurons, then it is considered that these neuronal responses are directly influencing the percept of the stimulus. This gives us a simple way to probe, non-invasively, the particular computational structure of the motion perception system. D. Tadin’s insights and results, including later work of collaborators and other researchers, are noteworthy because they showed how we can access the spatial and contrast characteristics of these motion tuned neurons.

The work of Schallmo et al. (2018) builds over the above very short overview about motion perception and its origin, particularly in the spatial and contrast domains through the use of D. Tadin’s experimental design. But they stated that the recent decade of surge in more application oriented studies, which are based on the accumulated knowledge from the earlier works on motion perception, is not totally justified because (citing their Introduction): “Strong assumptions are often made about the neuronal processes underlying these seemingly complex interactions between size and contrast during motion perception.”. Thus, they proposed that the overall effects and observations for the single grating design of D. Tadin and collaborators (and other researchers who use it) are not necessarily due to excitatory and inhibitory interactions between motion sensitive neurons but instead arises from the divisive normalization of neuronal activity, that emerges from classic neuronal network computations, and which they, and others, have used as a “computational principle”/”computational framework”.

In the following Research Advance manuscript, I comment on two methodological and theoretical aspects of their work, and demonstrate that the original design of D. Tadin does indeed allow to extract information about the putative excitatory and inhibitory processes that shape motion perception in the spatial and contrast domains. First, I discuss the psychophysics methods and analyses employed by the authors for the measures they provide. Second, I analyse their modelling approach and show that their model and its application contains conflicting knowledge with established computational neuroscience and behavioural modelling. From these analyses, in a last step, I develop the correct model for predicting motion perception in D.Tadin’s design and I show that such data can be used for *quantitative* inference of putative changes in underlying excitatory or inhibitory components, changes that partly can and partly cannot be inferred from the study of Schallmo et al. (2018).

## Results

The article of Schallmo et al. (2018) used five techniques for approaching the question about how contrast and size of the stimulus affect the necessary stimulus duration time to perceive the correct direction of motion. From these five methods, the psychophysics measures and modelling are the main foundations. These two methods are strongly intertwined in the argumentation and inferences that can be made from the design in the study. In the following, first two points about psychophysics and data processing are emphasized. Then, shortcomings in the modelling are discussed and the correct model is presented, fit on previously published data, and used to discuss the results of Schallmo et al. (2018).

### Psychophysics

#### Definition of the psychometric function in D. Tadin’s design

The method used by the authors is as follows. In this design observers were presented with a single moving stimulus whose contrast, size, presentation duration and direction of motion is varied across trials. The idea is to extract the duration threshold for various stimulus sizes and contrasts. Thus, observers are generally instructed (or learn it during the training before data collection) that (1) on each trial only one of two opposite motion directions can appear (e.g. left-or rightward directions), (2) on some trials the motion may appear so weak that it is not clearly perceived and they may need to guess its direction, and (3) they had to report the direction of motion they thought the presented stimulus had on the trial. These are necessary conditions for measuring the psychometric function in this design, and additionally researchers may, or may not, add more randomness from observer’s point of view by measuring thresholds for different contrasts and sizes of the stimulus within the same block. This design is a common “one-stimulus presentation two-alternatives forced choice” design, let’s call it here 1stim2AFC (Morgan, Watamaniuk and McKee (2000) called it Method of Single Stimulus; Kingdom and Prins (2009) called it 1AFC), that can be used for either measuring discrimination thresholds or measuring misperception effects (e.g. motion repulsion, Tzvetanov and Womelsdorf (2008); Tzvetanov (2012); e.g. tilt illusion, Westheimer (1990); Kapadia, Westheimer and Gilbert (2000)). In this particular design, the experimentalist is interested in the discrimination thresholds of the observers, defined as the necessary duration time of the stimulus for discriminating at some predefined level of correctness (let’s say 84%) the direction of motion of the stimulus. It is defined with respect to the midpoint of the psychometric function, as described below. The duration of the stimulus is varied between long durations (persons clearly see the motion direction) and short durations (persons have hard time deciding about the direction). The psychometric function spans the two dimensions of stimulus duration (continuous) and direction (binary). Therefore, the resulting data must be analysed as a single psychometric function spanning the full range of left-to-rightward motions (sign of the stimulus) of different durations (intensity of the stimulus). If we assign negative/positive as leftward/rightward motions, absolute intensity as the duration, and percent “rightward” responses as the y-variable one obtains a single psychometric function spanning the full range of 0 to 100% of proportion responses (Figure 2A). This definition clearly allows to define also the biases that can appear in this 1stim2AFC design, biases that can be due to response bias in insecure trials (decision to respond with always the same key when not sure) or perceptual biases (persons really see something different from “0”). Figure 2A depicts such two functions. The discrimination threshold is thus *x*(*p*=84%)-*x*(*p*=50%).

**Figure 2:**
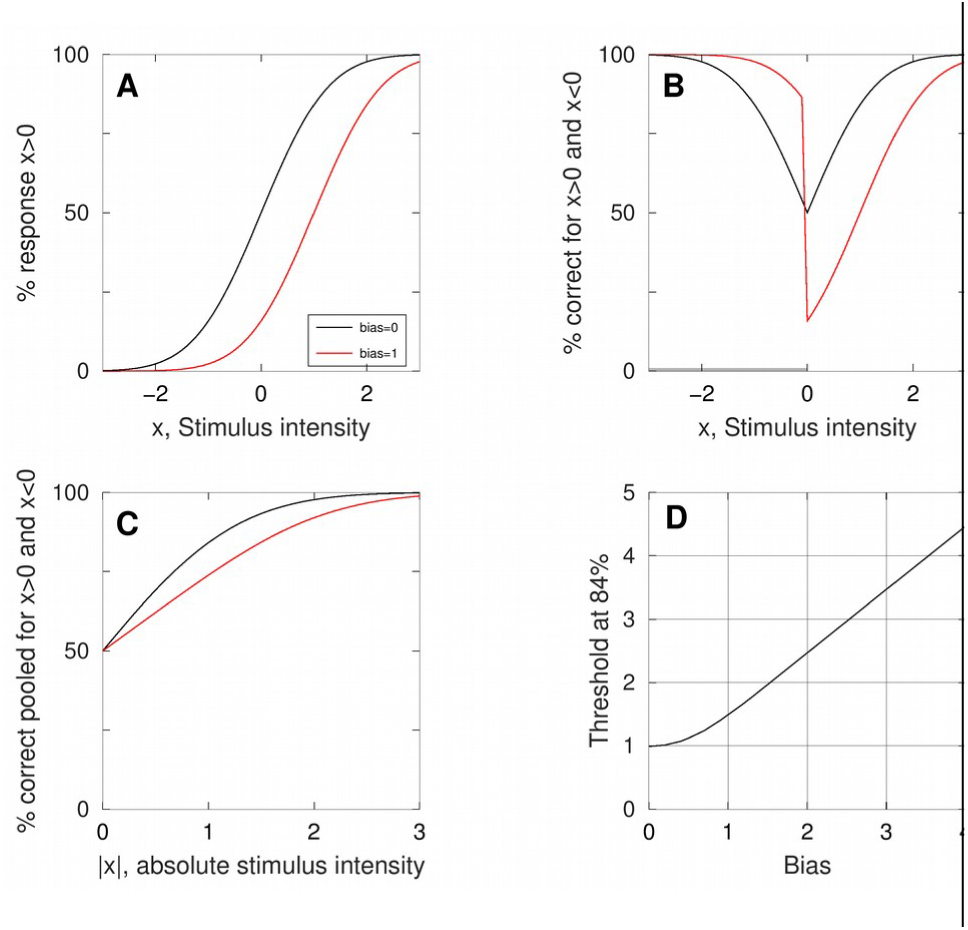
Psychometric function definition. (A) cumulative Gaussian function used to describe psychometric performance in 1stim2AFC designs; black and red curves plot no-bias and biased cases. (B) Replotting the function from (A) by defining Percent correct as the y-variable. (C) Pooling the negative and positive intensities into a single psychometric function of percent correct defined between 50% and 100% correct responses. (D) Threshold at 84% as a function of the bias from pooled psychometric function in (C). All plots have psychometric functions with standard deviation of 1.

Instead of analysing the data in this manner, Schallmo et al. (2018) defined the pooled psychometric function by computing the percent correct responses for each physical direction of motion (Figure 2B) and then pooled the data over the two directions (Figure 2C). The consequence of doing this is that, if a bias is present, this psychometric function does not define unambiguously the perceptual threshold of the observers, but also includes the bias. When one compares the extrapolated threshold in Figure 2C for the black (no bias) and red (bias of 1) curves, the change is ~52% higher. One can compute the threshold as a function of the amount of bias when no change in the real discrimination threshold is present (Figure 2D), demonstrating the strong change in thresholds with change in bias.

While one might argue that the results of the experiments still show strong and significant effects across conditions, which is undoubted, the reported thresholds have an unnecessary bias that is easily taken care in at least one of two ways: either through the data analysis procedure described above, or either at the experimental methods level by providing feedback to the observers about their response correctness on each trial. The later method was used by Tadin and Lappin (2005) while Schallmo et al. (2018) did not report to use it. This data analysis is important when one has to make model-to-data adjustment. (further details on psychometric function definition and plausible consequences on threshold estimation are presented in **Appendix A1**).

#### The composite variable “Size Index” involves simultaneously spatial suppression and summation

The authors reported their data by showing the mean duration thresholds and used the “Size Index” (SI) variable to indicate changes in spatial suppression and facilitation. They defined this variable as the difference of log-thresholds for the smallest and a larger size conditions at a given contrast level. In order for this composite variable to be representative of the concepts of suppression, which they defined as SI<0, and facilitation, which they defined as SI>0, it means that changes of thresholds between a small and larger size automatically labels the change as due to only suppression or only summation. This happens when: (1) at very low contrasts inhibition is very weak and thus the results are mainly influenced by spatial summation instantiated by excitatory mechanisms, therefore one should expect a decreasing threshold as a function of stimulus size (see Figure 1A, c=9%); and (2) at very high contrast inhibition is predominant and thus the results are mainly influenced by suppression, therefore one should expect an increasing threshold as a function of size (see Figure 1A, c=92%, but see the two data points for the smallest size). The overall psychophysical data of Schallmo et al. (2018) does not comply with these rules, in particular the low contrast data, which has some “V” shape, indicating that both excitatory and inhibitory mechanisms seem to act simultaneously in setting observers perceptual thresholds for motion.

#### Conclusion from Psychophysics

The type of data obtained from their design must be fit with the 0-100% percent responses range psychometric function, for example a cumulative Gaussian (Figure 2A), and then extract the discrimination thresholds of the observers. Then, contrary to the reported composite “Size Index”, the threshold data should be used directly to infer something about changes of inhibition and excitation in the visual motion system of the observers. This last point will appear very clearly once the model is established and used to predict how inhibition and excitation, together, shape the thresholds.

### Modelling

#### A note on normalization of neuronal activity

Here, before presenting the exact model for their experimental design, I want to comment on the issue of the computational principle/framework that the authors put forward. Schallmo et al. (2018) proposed “divisive normalization” as the binding and main framework for understanding facilitative and suppressive effects in motion perception, as other authors have proposed it for neurophysiology (Reynolds & Heeger, 2009; Carandini & Heeger, 2012).

How does the normalisation property appear? Let’s look at a classic neuronal network with excitatory and inhibitory nodes, respectively noted *y_e_* and *y_i_*. The activity of the excitatory node can be written:

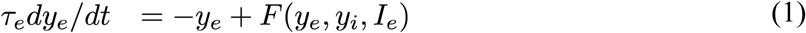

Here, *F*() is some function that describes how the nodes are connected and influence each other (through some weights, “firing rate”/”transducer” function, and the inputs to each node *I_e_* and *I_i_*) (Ermentrout, 1998). One general result for neuronal networks is that after a short time of activity the network stabilises itself on a “steady state” (but it could be also a pure oscillatory activity, for a simple introduction to the topic, see Wilson (2005)). What this means is that the activity of a neuron has an initial strong dynamics that typically shows oscillation activity just after input onset (e.g. a burst of firing), and then rapidly stabilises on some steady state activity (for model examples, e.g. Piëch, Li, Reeke and Gilbert (2013), or Wilson (2005)). This steady state activity can (rare) or cannot (in general) be computed in a simple analytical manner, depending on the exact functional form of the above network equations (the function *F*()). This stable point is what Schallmo et al. (2018) consider as a computational principle/framework (see also Appendix of Heeger (1992)).

From the above short presentation, it comes natural that neuronal activity stabilisation, aka normalisation, sometimes appearing as “divisive normalization”, accounts for the neurophysiological results, but it does so due to the exact functional form of the equations (transducer, excitatory and inhibitory effects…) that the modeller decided to put in it for matching the model to the data. They are carefully considered based on prior knowledge, especially neurophysiological but also behavioural reports. Schallmo et al. (2018), themselves, do fix their normalised activities by making the assumption that excitatory (facilitation) effects are in the numerator and inhibitory (suppressive) effects are in the denominator (their Equation 1 reproduced here):

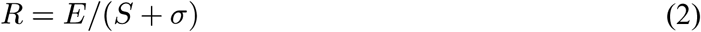

The exact reason for having such a form of stabilised activity is not given, out of the “divisive normalization principle” (although their model description hints to a classic feedforward model). This is only one possibility of normalisation result. The following two examples of neuronal networks will give further possibilities.

The first example is a V1-layer 4 cell from the model proposed by Grossberg and Raizada (2000) (their equations 17-18) for explaining contrast and attentional effects observed in neurophysiology of V1. The activity *y_ijk_* of these V1-layer 4 cells at equilibrium is:

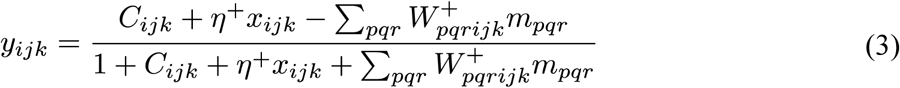

where *C_ijk_* are processed LGN inputs that are modulated by layer 6 cells activities (*x_ijk_*)organised in an excitatory-centre (*η^+^x_ijk_*) and inhibitory-surround (*W^+^_pqrijk_m_pqr_*) structure, this later being transmitted through a layer 4 inhibitory network activities *m_pqr_* (see their equation 19). Leaving aside further considerations about their model, this example shows how nominator and denominator include excitatory and inhibitory effects from various origins. It should be noted that these other feedback activities are themselves function of layer 4’s activities and other V1 layers, thus making the exact computation not analytically straightforward and necessitating numerical simulations.

The second example is a model of centre-surround interactions in motion processing taken from Kim and Wilson (1997) (their equation 6). The time dependent activity, *C_θ_*, of one neuron in a centre population sensitive to direction of motion *θ* is:

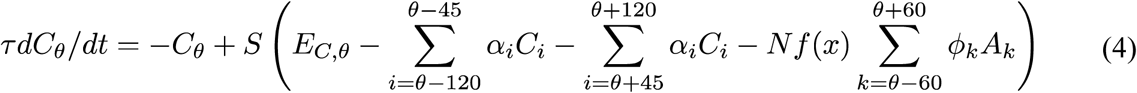

that gives at equilibrium the result:

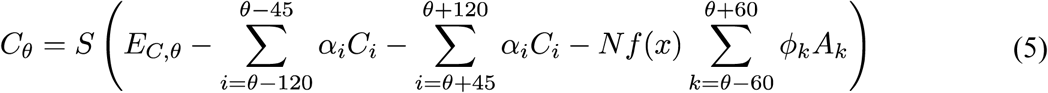

where *S*() is a simple transducer (*S*(*x*)=*x* if *x*>0, else 0), *E_C,θ_* is the preprocessed pattern motion input to the centre at direction *θ*, and *A_k_* are the activities of spatially surround neurons that themselves undergo excitation and inhibition in a similar manner (the remaining factors are network connection weights, see their tables 1 and 2). In this example the stabilized activities of the neurons are a simple function of input and subtraction of the inhibitory parts coming from the other cells with stabilized activities. If one wants to see a “divisive normalisation” in this example, we can say that the denominator is equal to 1, which is due to the particular network structure taken in the article. The normalisation property here comes from the facts that the input *E_C,θ_* is itself limited between a minimum a maximum, there are only inhibitory interactions, and the transducer does not allow for negative activities (see equation 4). In this case too, because of the intertwined interactions through all spatial and motion direction lateral connections, the computation of the stable activities were obtained through numerical simulations. (It is noteworthy to mention that Tadin and Lappin (2005) use a simplified version of such a network, see their Table 1 and further below, section **The correct model for D. Tadin’s design**).

What did we see from these examples? First, that the exact relation for the neuronal activity at equilibrium is not necessarily an excitatory effect divided by an inhibitory effect; second, that the relation can be widely different depending on the exact functional network and connectivities that are assumed by the modeller. The fact is: equilibrium activities of a neuronal network can be obtained from divisive normalisation; it is a simple plausible computation; the exact terms in the denominator and nominator depend on the network structure the researchers are interested in, based on preliminary knowledge coming from neurophysiology or from behavioural results hinting to a particular structure.

Putting “divisive normalisation” on a pedestal of a principle/framework as such, without consideration of the network structure of interest giving the normalisation property, some researchers may advocate (Reynolds & Heeger, 2009; Carandini & Heeger, 2012), but it is not the unique possibility.

#### Concerns in Schallmo et al. (2018)’s model of motion perception and its application

In their work, Schallmo et al. (2018) proposed that the threshold of perception, the parameter that is reported, is modelled through a simple rule of C*riterion* divided by model’s *Response* written as (Equation 3 in their Methods):

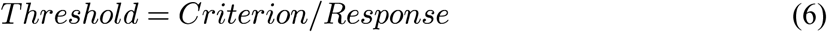

This equation is claimed to be somehow related to the DDM of decision making (Ratcliff, 1978; Ratcliff & McKoon, 2008; Forstmann, Ratcliff & Wagenmakers, 2016) (see also Link (1975); Link (1992); Smith (2000)), and it seems to have been taken from the two earlier studies of Tadin and Lappin (2005); Betts, Sekuler and Bennett (2012). The exact model description of how this equation is obtained is not fully described in Schallmo et al. (2018), neither in the two other studies using it (see **Appendix A2**), but instead refers to the study of Huk and Shadlen (2005). These later two authors investigated the detection of RDKs with behavioural and neurophysiological measures in monkeys on which they applied the DDM. But does the DDM predict a similar relation as in Equation 6? The interest in the drift-diffusion model is that it predicts the mean psychometric functions for proportion responses and decision times (response times) (e.g. Luce (1986); Palmer, Huk and Shadlen (2005)). The proposed equations are (Equations 4-5 directly taken from Huk and Shadlen (2005)):

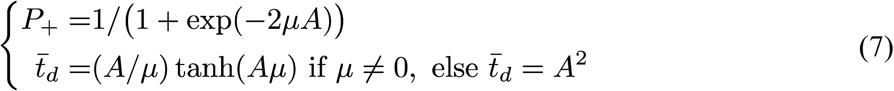

where *A* is the height of the boundary for decision making and *μ* is the drift rate. This later parameter is assumed to relate in a simple manner to model’s low-level response, *R*, for example *μ*(*x*)=*R*(*x*)×sign(*x*) (*x* is stimulus intensity, see **Psychophysics**). If we take the equation of *P_+_* and search for the *x_84_* value giving *P_+_*=*P_thr_*=0.84, one obtains:

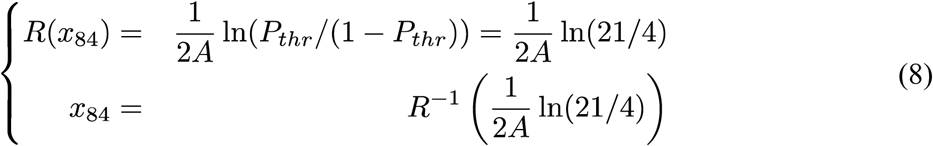

where the last equation is obtained by assuming that model’s low-level response *R*(*x*) is invertible. In the DDM result the stimulus *Threshold* is inversely related to the *boundary height A*, through some non-linear inverse function of model’s response. As such, Equation 6 is conflicting with the derivation obtained from the psychometric function of the DDM. The argument about using Equation 6 seems the fact that for a fixed *Criterion* stronger neuronal responses lead to shorter time *Thresholds*, and vice versa, thus providing a simple relation between neuronal responses and perception (see **Appendix A2**). Overall, their model relating perceptual *Threshold* and low-level model neuronal responses (Equation 6) is a new construct whose origin and derivation is not clearly given.

An important further concern in the model application done by Schallmo et al. (2018) is the way they left their model’s decision *Criterion* to differ between low and high contrast conditions (their Figure 2C,G and Appendix 1 – Table 1, first column of results). Because all conditions of size and contrast were presented randomly within the same block of measures (their page 12, Methods – Paradigm and data analysis – Psychophysics: “Staircases were run separately to determine thresholds for each of the six stimulus conditions (two contrasts x three sizes, as above). Condition order was randomized across trials.”) and there was no *prior* information about what will be presented on each trial, the decision criterion cannot differ between conditions measured within the same block. Allowing for different *Criteria* for different contrasts is not realistic because it means that the observers fixed their decision criterion on a trial basis *after* each stimulus was presented. Therefore, this is an issue that is conflicting with common ideas in detection theory.

#### General background for the modelling

Now we can turn to the modelling of the behavioural results in the simple experimental layout of D. Tadin. First let’s recall the factors of interest in such studies, {*size, contrast, duration*} of the stimulus, and the background hypothesis, that it is the excitatory-centre inhibitory-surround receptive field organisation in area MT/hMT+, which has a typical structure such that similar-to-centre motions in the surround reduce the activity due to centre stimulus presentation (Born & Bradley, 2005), that are a direct reason of the perceptual results obtain with motion stimuli.

In predicting psychophysical performance, the models are generally split into two independent stages, the low-level neuronal activities and the high-level decision stage. When combined they must predict the psychometric functions for each combination of size and contrast (see **Psychophysics**). The low-level activities are fed to the decision stage that only does one computation – to predict the dependent variable, here percent “rightward” responses.

An important point must be clarified concerning this experimental design and its modelling: the independent variable of the psychometric function is stimulus duration (*t_stim_*). Because of this, there is a very strong assumption in the modelling that is made: the neuronal activity, *R*, of the motion tuned neurons are a direct measure of the duration of the stimulus, exactly as for stimulus contrast. That is, there is some monotonic relation between stimulus duration and *R*, the low-level model of neuronal firing rate. This point is a necessary condition for being able to model the psychometric function that is assumed monotonically increasing with stimulus duration. Let’s assume that it has a classic saturating behaviour defined by a hyperbolic ratio equation (Albrecht & Hamilton, 1982) in the time domain:

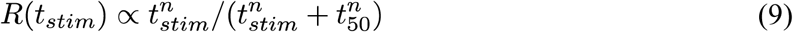

with *n,t_50_*>0. This equation states that for an infinitely long presentation of the stimulus the neuronal activity is finite. Furthermore, here it is assumed that this component is multiplicative of the “interaction” components.

Concerning the decision stage model, the two most common, and currently largely advertised and used, models will be considered. The first one is the classic Signal Detection Theory (SDT) (Green & Swets, 1966; Macmillan & Creelman, 2005) applied on neuronal activities (e.g. Britten, Shadlen, Newsome and Movshon (1992)). The second one is the “Drift-Diffusion Model” (DDM) (Link, 1975; Ratcliff, 1978; Luce, 1986; Link, 1992; Smith, 2000; Smith & Ratcliff, 2004; Huk & Shadlen, 2005; Palmer et al., 2005; Ratcliff & McKoon, 2008; Forstmann et al., 2016) that Schallmo et al. (2018) associate to their decision stage.

#### The correct model for D. Tadin’s design

After this long but necessary *aparté scientifique*, it is now possible to present a correct model predicting motion perception in D. Tadin’s design. We take the low-level processing stage of the model to be represented by a population of neurons coding the two possible motions with opposite directions, whose activity *R’*s depend on all independent variables of stimulus contrast (*c*), size (*s*), and duration (*t_stim_*). Here, both low-level models of Tadin and Lappin (2005) (but modified for having positive defined activities) and Schallmo et al. (2018), and both decision models (SDT and DDM) are considered, giving as model equation:

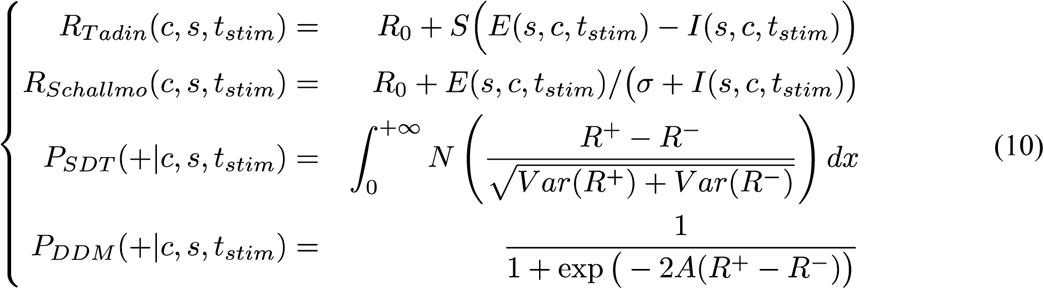

with *N*(*M*) representing a normal distribution with mean *M* and unit variance, and *R^+^* and *R^-^* representing the activity of neurons coding right- and leftward motion directions, respectively; *E*() and *I*() are the excitatory and inhibitory drives; *R_0_* is a “spontaneous firing rate” of the neurons (no input: *c*=0, or *s*=0, or *t_stim_*=0; but see **Discussion**); *S*() is the simple transducer from Equations 4-5; “+” sign in the psychometric function is defined as the rightward motions (see Figure 2); because the experimental design is a 1stim2AFC task with only two possible motion directions (see **Psychophysics**), SDT’s prediction is based on the difference of activities of the two motion coding populations, and *Var*(*R*) is assumed equal to *R* (Fano factor of one); for the same reason the drift rate *μ* in the DDM (Equation 7) is taken as the difference between the activities of neurons with opposite motion directions.

Important note: in the above Equation 10, activities *R*s of the motion tuned neurons may represent the equilibrium state values for the given stimulus input parameters, as discussed in section **A note on normalization of neuronal activity**, or more generally a mean activity across some time window of post-stimulus presentation; starting from equation 10 model presentation can be considered as following the tradition of “pattern analyzers” (Graham, 2011), where it is usual to directly use static mathematical models with predefined pattern sensitive inputs.

What remains to be defined are the excitatory and inhibitory drives *E*() and *I*(). For the moment we keep a general formulation. Because we assumed that neuronal response to stimulus duration, *R_dur_*(*t_stim_*), is independently pooled from responses to contrast and size, and that it has the same modulation on both components, we can write:

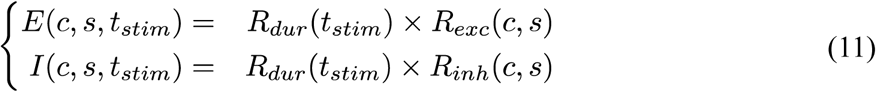

By using the threshold definition, 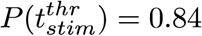, one finds for each low-level model considered here that (see **Appendix A3**):

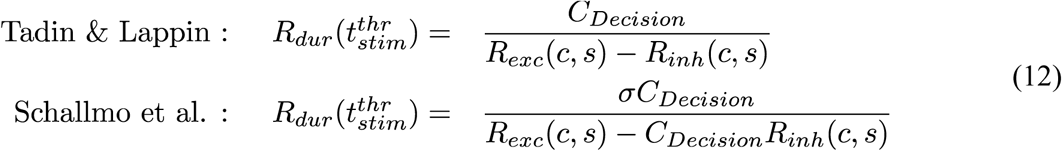

The constants *C_Decision_* are dependent on the decision stage models of psychometric functions as follows:

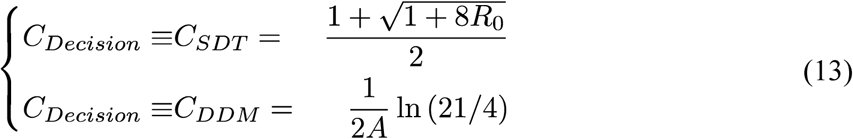

But these constants are NOT the criteria for giving the response, neither some “*arbitrary response value that must be reached to make the perceptual judgment*” (see **Appendix A2**). They are simply mathematical constants that are obtained from the full model assumptions and derivations, and what is defined as threshold for percent responses on the psychometric function (here 84%). In SDT case, the observer decision criterion is: if *R^+^-R^-^*>0 then give response “+”, else response “-”. The case of the DDM was presented in the previous section and it does contain the “boundary height *A*”, but in that case the decision criterion is which of the two boundaries is reached (if “+*A*” then give response “+”, if “-*A*” give response “-”; for further details see the DDM articles and reviews cited earlier), and the decision “height” is in the denominator.

Making the last step and computing the duration threshold of the theoretical observer, by using equations 9 and 12, one obtains:

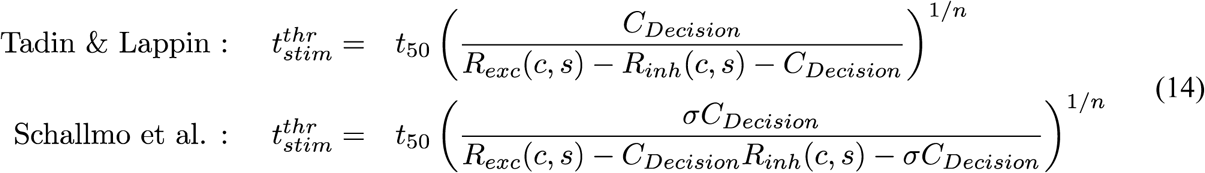

Therefore, although equation 6 proposed by Schallmo et al. (2018) has some similar form with the correct equation 14, its origin and interpretation are not satisfying. The correct general equations for modelling the psychometric functions of proportion responses as a function of stimulus parameters are given in equations 10, which from considerations about experimental threshold and model parametrisation *leads* to equation 14.

#### Comparing predictions of the two low-level models

Now that the derivations of the perceptual thresholds have been laid down, we can analyse whether these equations make interesting inferences about a particular model, or point to changes in thresholds as a function of excitatory or inhibitory strengths. Because the decision stage of the models make predictions that differ only in the constant *C_Decision_*, from now on, any application of a full model will be done with SDT. Furthermore, we fix, without loss of generality, in Schallmo et al.’s model *σ*=1 (amplitude rescaling of *E*() and *I*() in Equation 10).

The two models give the predictions of duration threshold with equations 14. Interestingly, the condition *C_SDT_*=1, which happens when *R_0_*=0, and assuming that *R_inh_*(Tadin)=*R_inh_*(Schallmo) and *R_exc_*(Tadin)=*R_exc_*(Schallmo), gives exactly the same final prediction from both models, independent from the particular functional form of the excitatory and inhibitory drives. That is, despite the very different initial model assumptions (Equations 10), one finds that the final prediction of threshold variation as a function of stimulus size and contrast are equal.

When *R_0_*>0, the exact difference between the two predicted thresholds depends on all three variables of excitatory, inhibitory and spontaneous firing rates. Nevertheless, even when *R_0_*>0, one can see that if the two models differ only in the inhibitory responses *R_inh_*, when *R_inh_*(Tadin)=*C_SDT_R_inh_*(Schallmo) both models still give exactly the same prediction of thresholds through a simple scaling of the inhibitory drives. Therefore, if the functions for excitatory and inhibitory effects in the two low-level models are chosen to be of the same mathematical form, then the two models can predict exactly the same thresholds by a proper rescaling of inhibitory amplitude.

To conclude, selecting either of the low-level models in Equation 10 is irrelevant for the final predictions. Consequently, it is the mathematical forms of the excitatory and inhibitory drives that predict the perceptual thresholds, not the “divisive normalisation” of Schallmo et al., neither the simple subtraction of Tadin & Lappin.

#### Application of the model

Until now, without taking a particular mathematical form of the contrast and size tuning properties, it was possible to make model analyses based on general considerations about the functions. Because the model predicts exact thresholds, and the deduced final equation 14 can be fit to the data (similarly to the previous applications of the resembling equation 6, see Tadin and Lappin (2005); Betts et al. (2012)), now we have to choose specific mathematical forms for the different components. By using Tadin & Lappin’s low-level model, based on prior knowledge, and writing the full equation of the model, we have:

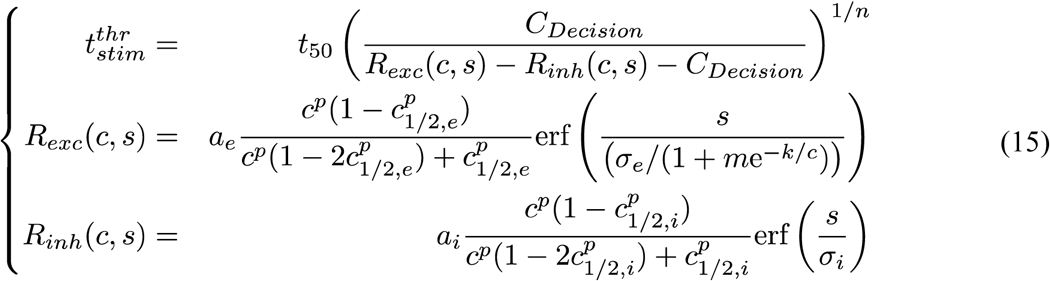

where {*a_e_,a_i_*} are the absolute response amplitudes of the excitatory and inhibitory components, the middle, complex-looking, ratio of contrasts in *R_exc_*() and *R_inh_*() is the classic hyperbolic ratio for contrast response functions (CRF) (Albrecht & Hamilton, 1982) (but rewritten such that at *c*=1 its response is 1 and at *c*=*c_1/2_* its response is 1/2), and the last part is the error function as a function of size (for the inhibitory part) and size and contrast (for the excitatory part) (Tadin & Lappin, 2005) (see also **Appendix A4** explaining why in this model excitatory-centre tuning width must vary with contrast). In the above equations it is assumed that the pure contrast responses for both excitatory and inhibitory drives have the same exponent *p* but different half-amplitude constants {*c_1/2,e_, c_1/2,i_*}. The size function, *erf*(), assumes that both receptive field components have a circular Gaussian shape and the space information is simply pooled across the stimulated area (stimulus centred on the receptive field). The model contains a total of 12 parameters: {*t_50_, n, C_Decision_, a_e_, a_i_, p, c_1/2,e_, c_1/2,i_, σ_e_, σ_i_, m, k*}. Without loss of generality, *n* and *a_e_* are set to 1 (parameters {*C_Decision_, a_i_*} are interpreted as proportional to the excitatory amplitude *a_e_*). This leaves a total of 10 free parameters.

##### Example of model fit

A demonstration of fit to the data of Tadin and Lappin (2005) is carried. There are enough data points (32) on a sufficiently large range of contrasts and sizes for constraining all 10 parameters. Fit results are: {*t_50_*=197, *C_Decision_*=0.051, *a_i_*=0.876, *p*=0.75, *c_1/2,e_*=0.0044, *c_1/2,i_*=0.081, *σ_e_*=2.5, *σ_i_*=3.6, *m*=7.78, *k*=0.305} (see continuous curves in Figure 1A and compare to the data). Although some parameters seem to have stronger influence on a single input dimension (e.g. *σ_i_* on size only), it is the 3D data, *Duration Threshold* vs {*Contrast, Size*}, that constrain all parameters simultaneously through the final non-linear model of Equation 15. Some interesting *quantitative* points can be derived from the final parameters: (1) inhibitory amplitude is quite strong, ~90% from the excitatory amplitude, which is a nice counterpart of neurophysiological results showing that short duration motion stimuli seem to preferentially stimulate neurons with strong surround inhibition (Churan, Khawaja, Tsui & Pack, 2008); (2) the constant *C_Decision_*=0.051 gives, if *a_e_*=100, *R_0_*~10 which is about 10% of *a_e_*, a rather common spontaneous firing rate of neurons (but see **Discussion**); (3) the excitatory half-amplitude constant *c_1/2,e_* is low, an interesting counterpart of neurophysiological observations for similar centre-surround designs showing low semi-saturation constants (Tsui & Pack, 2011); (4) the power of the contrast response function is very low (*p*=0.75), which was also reported to be the case in neuronal fits of size tuning data of MT (Tsui & Pack, 2011).

In summary, the correct computational model fit to the behavioural data provides *quantitative* parameters that globally match expectations about parameter values gathered from neurophysiological studies. Thus, one can apply the model on behavioural data to infer putative changes of inhibitory or excitatory mechanisms under experimental manipulations, experimental manipulations that were performed by Schallmo et al. (2018) (although the data may contain some biases, see **Psychophysics**, here it is assumed that these biases correspond to a constant factor across all thresholds, thus simply scaling them all; this is done in the model by changing the global time constant, *t_50_*).

##### The psychophysics measures of Schallmo et al. (2018)

In their study, the authors reported duration thresholds for a total of six different experimental conditions, 3 sizes and 2 contrasts. Therefore, the data does not allow any direct and exact model fit. Despite this, given the above prior knowledge and tests, we can make the assumption, based on the hypothesis of same origin of effects, that parameters should be globally similar between the different studies; especially here the excitatory part. Thus, we fix: *c_1/2,e_*=0.005; *p*=0.75; *C_Decision_*=0.05; *σ_e_*=2.5; *m*=7.8 and *k*=0.3; parameter *t_50_* is left to vary because it simply scales the overall data (on a log-scale it shifts the curves up or down). Thus, the interesting parameters about inhibition are left free: amplitude (*a_i_*); half-amplitude constant (*c_1/2,i_*), and inhibition tuning width (*σ_i_*). A fit gave: *t_50_*=421, *a_i_*=0.74, *σ_i_*=7.7, *c_1/2,i_*=0.072; the results are depicted on Figure 3A. It shows that the data are globally explained, although some discrepancies at medium size stimuli and low contrast are present (this discrepancy hints to some model shortcomings, see **Appendix A5**).

**Figure 3:**
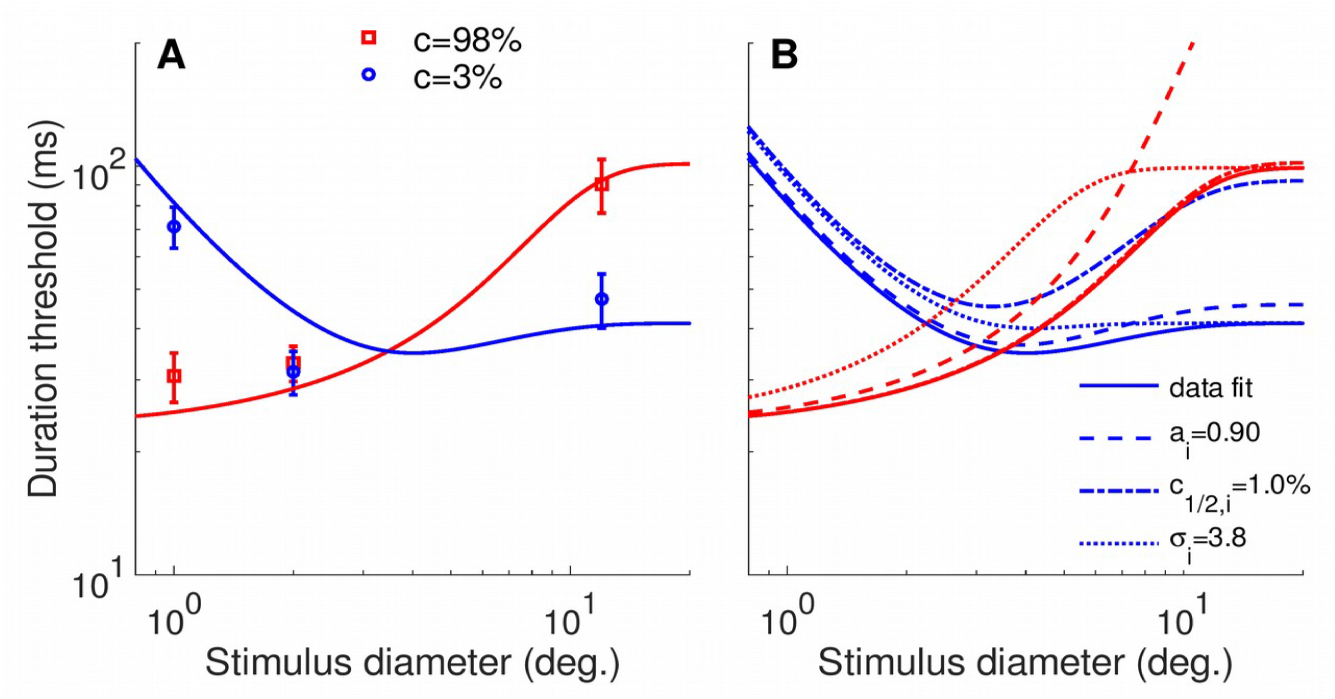
(A) Fit of model to the data of Schallmo et al. (2018), data from their Figure 2C. (B) Model prediction of threshold changes when only one among the three parameters (*a_i_, c_1/2,i_, σ_i_*) is changed such that inhibition is increased. “Data fit” curves are from (A). Red curves are for high contrast stimuli and blue curves are for low contrast stimuli.

##### Manipulating individual observer’s level of inhibition

What should we expect as an outcome in this experimental design if the overall level of inhibition within a person is manipulated? In the model the effects of the inhibitory component are instantiated through three model functions, one of amplitude, one of contrast, and one of size, with their respective parameters {*a_i_, c_1/2,i_, σ_i_*}. The effect of enhanced inhibition can affect all three functions. Thus, there are three simple ways to obtain stronger inhibition: an increase in amplitude (*a_i)_*), a decrease in contrast half-amplitude constant (*c_1/2,i_*) such that inhibition is activated at lower contrasts, and a decrease in tuning width (*σ_i_*) such that inhibition is activated earlier in spatial domain. These three simple examples are contrasted in Figure 3B to the results of the previous data fit (Figure 3A). As one can see, amplitude increase affects mainly thresholds at large stimulus sizes, half-amplitude decrease mainly affects low contrast responses and thus changes thresholds at low contrast, and inhibitory size decrease affects thresholds for small to medium size stimuli. One important result is that all three changes *increase* duration thresholds. This can easily be understood by referring to equation 15, where one can see that increasing inhibition (*R_inh_*(*c,s*)) deceases the denominator which in turn increases the overall bracket. That is, while the exact threshold variation is dependent on the combined changes of inhibitory parameters, increase in inhibition makes overall thresholds higher.

The experiment of Schallmo et al. (2018) about “**Pharmacologically enhanced inhibition**” indeed show such a behaviour. That is, duration thresholds of the drug-intake condition increased when compared to the placebo condition. Thus, contrary to their conclusion from the results (p. 7: “This result is not consistent with the idea that spatial suppression is directly mediated by neuronal inhibition.”), their results are consistent with the original hypothesis of drug-based inhibition enhancement of spatial suppression (if one assumes all other components are unchanged, see **Discussion**).

##### Comparing inhibition parameters between observers

In a last interesting experiment, the authors compared duration thresholds and non-invasive measures of GABA+ concentrations in visual cortices. They found that mean individual thresholds anti-correlated with the concentration of GABA+, obtained from Magnetic Resonance Spectroscopy, of the observer. This, as they stated it, is rather interesting because, as it was shown just above, stronger inhibition increases duration thresholds. Nevertheless, the exact contribution of the inhibitory component to the mean individual thresholds is unknown because the individual thresholds are dependent on all factors of excitatory and inhibitory drives, plus the individual’s half-amplitude time constant (*t_50_*, see Equations 9 and 15) and observers’ spontaneous neuronal activity. In order to know each component of each observer, a full model approach is necessary with the corresponding measures.

#### Conclusion from modelling

From the above overall description few conclusions are drawn. First, the computational model predicting the duration thresholds in D. Tadin’s design is simple and easy to obtain. From careful considerations about its basic assumptions and functional form of the neuronal activities, it was shown that two “competing” models of neuronal activities, i.e. “divisive normalization” or simple subtraction of excitatory and inhibitory drives, cannot be dissociated in this design. Indeed, by an amplitude change of inhibitory component between the two models they allow to predict exactly the same thresholds. Thus, it is the exact mathematical form of the excitatory and inhibitory drives that defines the shape of suppressive and facilitative effects on duration thresholds as a function of size and contrast. Second, the computational model was shown to provide quantitative estimates of parameters that globally matched known neurophysiological counterparts. Third, the application of the model to the experimental design of Schallmo et al. (2018) showed (1) that in the experiment with pharmacologically manipulated inhibition the behavioural results of increased thresholds are consistent with stronger inhibition, and (2) that to know exactly the amount of inhibitory change under a given manipulation or between different observers it is necessary to have a full model approach.

## Discussion

This Research Advance work analysed the spatial summation and suppression effects that raise in human perception of visual motion, specifically in the very simple stimulus design of D. Tadin (Tadin et al., 2003). It was prompted from the report of Schallmo et al. (2018) that (1) “divisive normalization” explains this paradigm’s results instead of more classic considerations about excitatory and inhibitory components, and (2) that their results seemed to show that suppression is not driven by inhibitory mechanisms. Therefore, this work presented the model for predicting the perceptual thresholds in the experimental design of D. Tadin and made multiple inferences from it. The conclusion from the model were presented in the relevant section. Here are discussed some points about psychophysics and modelling that were not developed further in the **Results** section.

When testing computational models of perception, we match the model prediction to the behavioural measures obtained in a particular experimental design through the use of the psychometric function (PF). Its exact form, and thus definition, depends on the experimental design. In the “1-stimulus-2-Alternatives-Forced-Choice” (1stim2AFC) design analysed here, it was argued that the PF should be defined as a function of motion duration and direction, thus representing a typical PF for “discrimination” (discriminating between two possible motion directions) that can be defined in the full range of percent responses (0% to 100%). Because in this paradigm PF thresholds (inverse of slope) vary in a large range, researchers customary report results in pooling opposite motion directions that allows to plot the results on log-scale of stimulus intensity. It provides a simple visualisation of the threshold change that is seen as a shift of the PF on the log-scale data (this representation is similar to the classic “detection” results representation, e.g. typical contrast “detection” in 2stim2AFC designs). The current work allowed to model the psychometric function by considering the inputs and outputs that the model should incorporate and predict. It provided the general model equation (Equation 10), where the low-level model responses are presented together with the decision stage method, that led to predict the perceptual thresholds.

The model approach in this work is based on “static model”, or “pattern analyzers” (Graham, 2011). It approximates the responses of the neuronal populations to the inputs in specific ways by assuming that all neuronal interactions lead to simple “static” functional forms.

First, the major independent variable stimulus duration, *t_stim_*, was modelled as activating neurons independently from the two other variables of stimulus contrast and size. The results show that it is a sufficient assumption. Furthermore, it was hypothesized that the exact way this variable affects neuronal responses is to modulate independently both excitatory and inhibitory drives. On the contrary, one can hypothesize that the effect is to modulate the combined response of excitatory and inhibitory components. While the “subtractive inhibition” model of Tadin and Lappin (2005) is not affected by this difference, the “divisive normalization” model of Schallmo et al. (2018) gives different predictions (see **Appendix A6**).

Second, the model incorporates the “spontaneous firing rate” of the neurons, *R_0_*, as the minimum possible activity. Nevertheless, spontaneous activity of the neurons are known to be lower for surround-suppressed cells in comparison to non-suppressed cells (Churan et al., 2008). This hints to the possibility that, when a moving stimulus is presented to them, the activity of the neurons sensitive to opposite motion directions is in fact lower than *R_0_*, an effect already reported (Snowden, Treue, Erickson & Andersen, 1991; Britten, Shadlen, Newsome & Movshon, 1993) and successfully modelled (Simoncelli & Heeger, 1998), and that it may vary with stimulus size and contrast. The model might be improved by including such an effect.

Third, it is known that the neuronal responses have contrast and size tuning such that at least one of the drives, excitatory or inhibitory, has intertwined contrast and size response functions. In the current work only the size tuning component of the excitatory drive was assumed to be changed with contrast variations. It is possible that instead of using size tuning to change with contrast, one can describe behavioural results by using the contrast tuning components to change with size variations. Such a possibility was used by Tsui and Pack (2011) for modelling their neurophysiological results and one might successfully apply such a model also to the behavioural results.

Last, if one manipulates inhibition, as performed by (Schallmo et al., 2018), the question naturally arises of how this experimental manipulation should be incorporated in the model. Here, it was argued only through the change of the inhibitory drive model. Since the model approach is based on “static” models, one cannot discard the argument that all the components of the model of “steady state” activity are modified. This is so because in changing connection weights of one type (excitatory or inhibitory) in the system affects the global equilibrium, and this later part is modelled, not the real weights. Therefore, because the experimental design of D. Tadin simultaneously measures the spatial and contrast domain characteristics of motion perception, the model necessarily must incorporate all these factors in its excitatory and inhibitory drives, which naturally leads to high multidimensional parameter space (here 10 parameters). The consequence is that, unless there is prior knowledge, the experimental measures must be carefully designed and carried in order to have data that can correctly constrain all parameters.

This work showed that we are very successful in providing simple and elegant modelling of behavioural results, results that demonstrate interesting perceptual phenomena that we associate to neurophysiological counterparts. But our modelling must be carefully considered and weighted with respect to prior knowledge. When neurophysiological, computational and behavioural results are combined in appropriate conditions for comparisons, we do gain important insights and knowledge about a given neurophysiological system and how it affects perception. In the case of suppression and facilitation of motion perception in humans, our understanding of how the percept is created and modulated has strong grounds from which we can make interesting inferences and, importantly, allow us to go toward more application oriented studies.

## Appendix

### Appendix A1

#### Psychometric function definition, fitting and threshold estimation

As presented in the main text, Schallmo et al. (2018) defined the psychometric function between 50% and 100% correct responses on stimulus intensity range -∞ to +∞. They used the common adhoc Weibull function of the form:

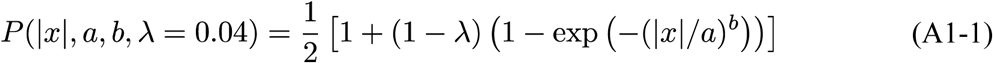

with the two parameters {*a,b*} defining the threshold (*a*) and slope (*b*), *λ* is observer’s lapse rate, and *x* is the stimulus intensity (negative values are “leftward” motions).

A first issue is that the above definition states that the psychometric function might have a sigmoidal form (if *b* is much bigger than 1, from ~2 and above) on a linear scale of *x* that starts from zero. What it means is that below some absolute intensity of the stimulus, |*x_low_*|, observer’s response is near chance level. This value *x_low_* depends on the “slope” parameter *b* of the Weibull function as well as the threshold *a*. The natural question coming from this definition is: If this effect exists, what is its origin? Because modelling of the psychometric function includes two levels, low- (stimulus processing) and high-level model (decision stage), there are two possibilities to consider. Either (1) this effect is based on some low-level sensory processing that must be carefully considered and incorporated in the model, either (2) this effect is a high-level decision stage process that should also somehow be incorporated.

It turns out that case (2) was already proposed and discussed for the experimental design of 1stim2AFC. Garcia-Perez and Alcala-Quintana (2013) proposed that for very weak stimuli observers may have an *indecision interval* (they quote Fechner as the first to mention this “interval of uncertainty”) where observers are not sure about having seen something relevant for deciding, and thus decide to guess. This allows to explain typical observations in the 1stim2AFC design of possible psychometric function shifts when observers always use the same response to respond in “undecided” trials. Because of the data processing method in Schallmo et al. (2018) (see main text) the above indecision model will shift the “Weibull” psychometric function toward higher stimulus intensities and steeper functions (higher slopes). Even in symmetric use of the two possible answers, when the interval of indecision is not zero it should create sigmoidal psychometric functions, with steeper slopes, than predicted by the simpler model of low-level + classic decision stage. Thus, it is possible that the ad-hoc Weibull function fits with high slopes *b* of the 1stim2AFC design do correspond to the model predicted data. Interestingly, because of their exeprimental methods (all stimuli presented within the same block), the above effect should be the same across all psychometric functions that were measured. Thus, one should expect that it corresponds to a scaling of the thresholds toward higher values, a scaling that is a simple shift on log-scale.

Overall, psychometric function fitting should be carried as discussed in section **Psychophysics** in order to avoid the above issues.

### Appendix A2

#### Relation between the model of Tadin and Lappin (2005); Betts et al. (2012); Schallmo et al. (2018) and the drift-diffusion model

The proposed model for predicting duration Thresholds (Equation 6) seems to be related in some loose way to the drift-diffusion model without clear description of the way it was obtained, out of the fact that this rule allows to relate inversely duration Thresholds to neuronal activity of the model (*R*). Here are quotations from their articles where the authors describe and discuss their model and its putative origin.

In Tadin and Lappin (2005), the authors state (page 2063):

> “Threshold (T) was taken to equal the number of times a response (R) needed to be repeated to reach a certain *Criterion—essentially modeling “how long” the response needed to be maintained in order to generate sufficient evidence about the motion direction of the stimulus* [emphasis by author TT]. This computation models the accumulation of evidence that likely underlies duration threshold measurements (see Footnote 1).”

And their Footnote 1 states:

> “Use of duration thresholds was based on the assumption that if the neuronal response to a stimulus is weak and/or noisy, then longer stimulus exposure will be required for correct perception. More specifically, deciding whether an object is moving in one of two possible directions can be conceptualized as a process involving accumulation of sensory evidence over time (Gold & Shadlen, 2000; Roitman & Shadlen, 2002). When neuronal responses are noisy or attenuated, as with a highly suppressed motion stimulus, sensory evidence accumulates more slowly and a correct decision thus may require longer exposure duration (Roitman & Shadlen, 2002).”

Betts et al. (2012) state (page 2):

> “The use of the stimulus duration threshold as a psychophysical measure of behavior is based on the idea that a perceptual decision regarding the direction of motion occurs only after a sufficient amount of information has been presented (i.e., after some response criterion has been surpassed) (Huk & Shadlen, 2005; Roitman & Shadlen, 2002). In this way, *weak and/or noisy signals require longer stimulus presentations to surpass the criterion for the perceptual response* [emphasis by author TT]. In the context of the model, the threshold, T, is the duration for which the response of strength R must be presented in order to surpass the Criterion (Tadin & Lappin, 2005). The *Criterion parameter determines the vertical placement of the model output on log–log coordinates, and inverts the independent variable such that increased responsiveness produces decreased duration thresholds* [emphasis by author TT].”

Last, Schallmo et al. (2018) state (page 13):

> “To equate the response predicted by the model to motion discrimination thresholds, we assumed an inverse relationship between predicted response and discrimination time, such that:
>
>
> 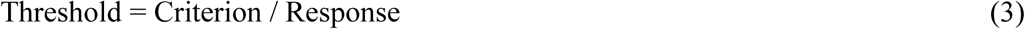
>
>
> where Threshold is the amount of time needed to discriminate the direction of stimulus motion, *Criterion represents an arbitrary response value that must be reached to make the perceptual judgment* [emphasis by author TT] (Huk and Shadlen, 2005), and Response is the predicted response rate (from Equation 1). This framework is consistent both with previous modeling efforts (Betts et al., 2012; Tadin and Lappin, 2005)[…]”

From the above snippets of their publications, it appears that (1) the decision stage model the authors advocate in their work has some loose relation to the DDM model and (2) this decision stage seems more a practical construct for predicting the dependent variable by the model.

Thus, despite tentatives to connect them, the comparison of DDM and their model that is carried in the main text shows that the two models are different.

### Appendix A3

#### Derivation of Equation 12

From Equation 10 we can write:

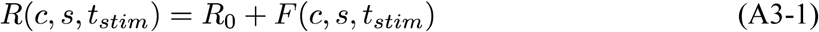

where *F*≡*F*(*c,s,t_stim_*) is the general non-specific function of the model. Then, using the SDT model, the definition of the threshold, and defining *F^+^*≡*F*+*R_0_* and *F^-^*=*R_0_*, we have:

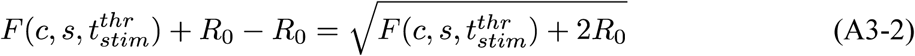

Which after simplification gives the second order equation and its solution:

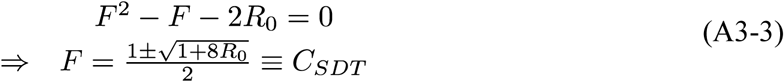

In this last equation the negative solution is not biologically meaningful and thus discarded. A similar to the last equation is obtained in the case of the DDM. From here, one derives the main text equations 12 and 14.

### Appendix A4

Here is analysed the question of what kind of excitatory and inhibitory drives can, or can not, predict the behavioural results. One noteworthy point from Equations 12 and 14 is that they allow simple and interesting inferences, inferences that, to the best knowledge of the author, for the particular design of D. Tadin, were missing until now.

#### Independence of contrast and size effects can not predict behavioural results: simple case

If we make the assumption of independence of stimulus contrast and size effects on each drive, *R_exc_*(*c,s*)=*R_c,exc_*(*c*)*R_s,exc_*(*s*) (similar equation for inhibition), and that inhibitory contrast response function, *R_c,inh_*(*c*), is a scaled version of the excitatory one, that is, *R_c,inh_*(*c*)=*kR_c,exc_*(*c*)≡*kR_c_*(*c*), for all models one obtains:

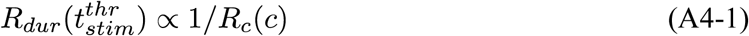

That is, because of the monotonic relation for each function on both sides of the above equation (both *R_dur_*(*t_stim_*) and *R*(*c*) are monotonically increasing with their variable), duration thresholds are inversely related to contrast for any size of the stimulus. In fact, the behavioural results for small stimulus sizes follow this relation (see Figure 1), but not for large stimulus sizes. Thus, independent effects of stimulus contrast and size cannot predict the psychophysical observations of inverted relation between duration thresholds and contrast at large stimulus sizes. We can discard this possibility as incompatible with behavioural evidence. This result is a nice counterpart of the neurophysiological findings in low-level visual processing systems, that stimulus contrast and size are somehow strongly intertwined in neuronal responses of areas V1 and MT (Sceniak, Ringach, Hawken & Shapley, 1999; Cavanaugh, Bair & Movshon, 2002; Tsui & Pack, 2011).

#### Independent contrast and size effects in the model: different, non-scaled, contrast tuning of excitatory and inhibitory components

Here is further analysed the condition of independent effect of contrast and size. Now it is assumed that *R_c,inh_*(*c*)≠*kR_c,exc_*(*c*), for example a simple shift of the inhibitory contrast response toward higher contrasts, and we assume that the components describing contrast responses are normalised such that their responses at *c*=1 is equal to one.

Then we can predict the inverted effect of contrast on duration thresholds as a function of stimulus size (e.g. taking the ratio of high-to-low contrasts of equation 14 one finds a simple relation between size and contrast functions and can deduce the relation at which the inversion happens; not shown). But the behavioural data show a very strong variation with contrast of the “minimum” (or dip) of the duration threshold as a function of stimulus size. In the data of Tadin and Lappin (2005) this minimum changes from ~2-3 degrees at c=9% down to ~0.5-0.6 degrees at contrasts 42% and 92% (see Figure 1A), a factor change of ~3-6. To demonstrate that this change is hardly predicted with independent contrast and size effects, we take for the size functions the error functions:

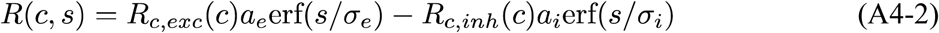

Because the dip of the effect happens when the response *R*(*c,s*) reaches a maximum on the size dimension (in Tadin & Lappin’s model; but in Scahllmo et al.’s there is also the decision constant), we can compute the stimulus size at which the dip appears, *s_dip_*, which gives:

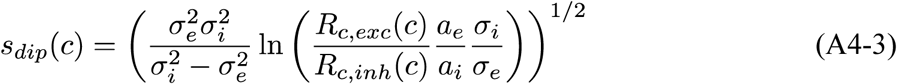

From this result, it is possible to find the necessary ratio of excitatory to inhibitory low-contrast responses for observing a given dip change as a function of the ratio of high-contrast responses. By using the ratio between size dips at low and high contrasts, *r_dip_*=*s_dip_*(*lc*)/*s_dip_*(*hc*) (*lc* – low contrast; *hc* – high contrast), it is found that:

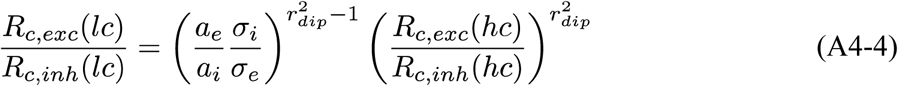

If we take the best case of *R_c,inh_*(*hc*)=*R_c,exc_*(*hc*), e.g. at contrast of one the normalised contrast coding components are equal, and *a_e_*=*a_i_* (maximum surround inhibition), then we have:

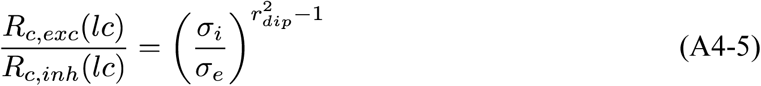

That is, the ratio of excitatory-to-inhibitory low-contrast responses is a function of the ratio inhibitory-to-excitatory size tunings to the power of squared *r_dip_* ! This model can predict factor of dip changes of ~3–6 only if the inhibitory responses at low contrasts are essentially zero (*R_c,inh_*(*c*)=(*σ_i_/σ_e_*)^-35^*R_c,exc_*(*c*) – (*σ_i_*/*σ_e_*)^-8^*R_c,exc_*(*c*)). This would correspond to a strongly expansive inhibitory contrast response function at intermediate to high contrasts, and thus explains the observation that this model does not correctly predict the behavioural data (Tadin & Lappin, 2005).

To conclude, the model of low-level responses cannot have independent contrast and size effects. Some form of contrast dependent size tuning widths is necessary to predict the behavioural results, as argued by Tadin and Lappin (2005) and reported neurophysiologically (Sceniak et al., 1999; Cavanaugh et al., 2002).

#### Necessity of centre excitatory tuning width to vary with contrast

One can further analyse which single component, excitatory or inhibitory, influences the most the position of the dip. We can again use Equation A4-3 and rewrite it as:

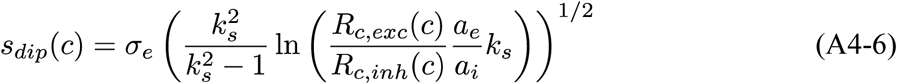

where *k_s_*=*σ_i_/σ_e_* is the ratio of inhibitory-to-excitatory size tuning. From this equation it is much easier to see that, at fixed ratio *k_s_*, the dip position is linearly related to *σ_e_*, while at fixed *σ_e_* it is much slowly changing with *σ_i_*. Thus, the behavioural data of duration thresholds showing large changes of the dip position can only be explained if excitatory centre tuning width is contrast dependent. The other factors of “pure” contrast responses and inhibitory tuning widths depending on contrast do not allow for large changes in dip positions.

### Appendix A5

#### Measures of spatial suppression and summation with sine gratings do not conform to the model

The data from the RDKs of Tadin and Lappin (2005) presented in Figure 1 are well fit by the model. In an attempt to explain also other data, it was found that the model has difficulties in fitting them. These data were obtained with moving sine grating stimuli, that were seen through Gaussian envelopes (Gabor patches). The fitting was unsuccessful in matching the model to the data. When the 10 parameters were left free the final best fit gave biologically not plausible values; even with these best parameters, the fit was still difficult to reconcile with the data. The reason of this discrepancy can be seen in Figure A1. Panels A-C replots the data from three studies that used moving sine gratings at different contrasts and sizes. As it can be seen, the grating data has a particular behaviour: at low contrasts (~3%) the thresholds have the typical decrease with increasing stimulus size; at slightly higher contrasts (4.2-5.5%) the data show a U-shape with increasing stimulus size; from contrasts above 10% the thresholds exhibit mainly monotonic increasing behaviour with possible plateau effect at very high contrasts (panel C). Panel D presents the model predictions. It can be seen that the model shows a typical U-shape of thresholds vs. stimulus size for all but the lowest contrasts of 2.8-5.5%, where it is very weak. One can use Equation A3-6 to see that at very high contrasts one has approximately:

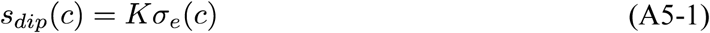

with *K* approximately a constant (*k_s_* increases with contrast and the ratio of contrast functions slowly decreases toward 1, which makes the bracket in Equation A4-6 to increase very slowly with increasing contrast, in opposite direction of *s_dip_* variation). Thus, one has the choices: (i) to change the function relating excitatory centre size and contrast such that at high contrasts it drops much rapidly as in the data, or instead (ii) suppose that the valid fit in Figure 1 hints to model inadequacy for explaining results with grating type of stimuli.

**Figure A1:**
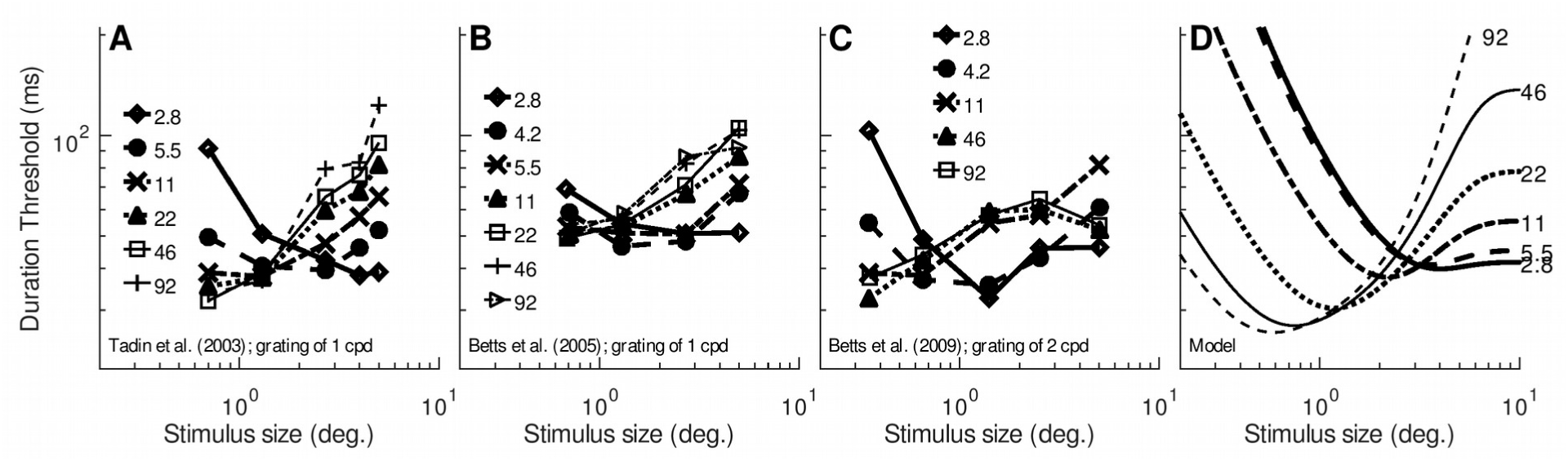
Duration thresholds data from reports using sine grating type of stimuli: (A) Gabor patches results from Figure 1 in Tadin et al. (2003), (B) Gabor patches results for young observers from figure 2 in Betts, Taylor, Sekuler and Bennett (2005), (C) Gabor patches results for young observers from Figure 1 in Betts, Sekuler and Bennett (2009); and (D) model (Equation 15) prediction for parameters {*t_50_*=400, *C_Decision_*=0.05, *a_i_*=0.9, *p*=0.75, *c_1/2,e_*=0.005, *c_1/2,i_*=0.08, *σ_e_*=2.5, *σ_i_*=4, *m*=8, *k*=0.3}. Legend in each panel for different contrast of stimuli. In (D) contrasts are printed on the rightmost end of each curve.

#### The data at low contrast of Schallmo et al. (2018)

The above short section showed that there are difficulties in matching the model to grating type of stimuli. In analysing Schallmo et al. (2018)’s data, it is apparent that their results for low contrast condition exhibited a different behaviour, a “U-shape” for thresholds vs. stimulus size; this U-shape is very pronounced in their Figures 4 and 5. This seems to match the U-shape results for slightly higher contrasts reported in the section above. Given that the stimuli of Schallmo et al. (2018) differ in the window type used (circular window with blurred edges), it is difficult to make comparison between the studies using moving gratings. Despite this, it should be noted that leaving the full model (10 parameters) to fit the 6 data points of Figure 2C in Schallmo et al. (2018) also gives excellent fit but with biologically not meaningful values (results not shown); a similar outcome as when trying to apply the model to Gabor patches stimuli.

Thus, it seems sine grating type of stimuli have peculiar results that are not explainable with the model derived in the main text.

### Appendix A6

#### Schallmo et al. (2018)’s model with global modulation by *R_dur_*(*t_stim_*)

In Equations 10-11, it was assumed that *R_dur_*(*t_stim_*) modulates independently each of the two drives. Instead, one can assume that *R_dur_*(*t_stim_*) modulates the full response to the combined drives. In this case, Schallmo et al. (2018)’s model provides slightly different equations, which are presented here.

The low-level response of the model becomes:

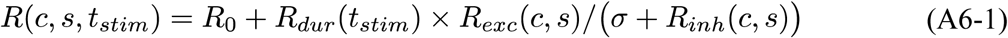

Remarque: one may think to re-interpret the above equation as: “only the excitatory drive is modulated by stimulus duration”; if one recalls that this is a “static” model of the final “equilibrium/steady state” activities, the interpretation should not be mistaken between the two possible implementations.

From equation A6-1 and the decision stage model, at threshold one obtains:

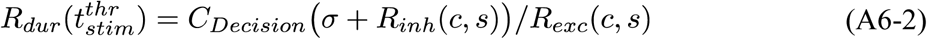

Which, by using Equation 9, gives:

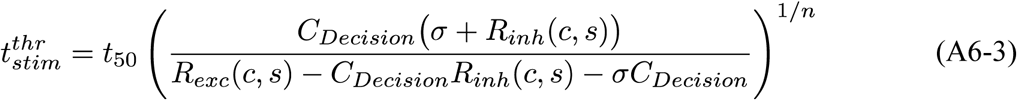

This last equation can be compared to the prediction of the main text (equation 14). This equation has a main effect to create inhibition effects on duration threshold much stronger than the main text model. This equation also provides a fit of the data of Tadin and Lappin (2005) (not shown). As for the main models (see **Appendix A5**) this model also succeeds to fit the Gabor patches results, but with biologically unrealistic parameters (not shown). No further analyses are provided here, neither detailed comparisons with the main text model.

